# ALG13 loss-of-function alters glycosylation, impairs neuronal maturation, and drives network hypoactivity in a cortical organoid model of CDG

**DOI:** 10.1101/2025.07.09.663964

**Authors:** Rameen Shah, Rohit Budhraja, Silvia Radenkovic, Graeme Preston, Alexia Tyler King, Sahar Sabry, Charlotte Bleukx, Ibrahim Shammas, Lyndsay Young, Jisha Chandran, Seul Kee Byeon, Ronald Hrstka, Doughlas Y Smith, Nathan P. Staff, Richard Drake, Steven A. Sloan, Akhilesh Pandey, Eva Morava, Tamas Kozicz

**Author notes:** Senior Author (Eva Morava, Tamas Kozicz). Equal contributions (Rohit Budhraja, Silvia Radenkovic, Eva Morava, Tamas Kozicz).

## Abstract

**Background:** Congenital disorders of glycosylation (CDGs) are a group of rare metabolic diseases recognized for their neurological presentations, including developmental delay and seizures. However, the link between glycosylation defects and cortical brain network pathology remains elusive.

**Methods:** To address this unmet need, we generated iPSC derived human cortical organoids (hCOs) for ALG13-CDG, which is the second most common CDG that is also X-linked. To comprehensively understand the impact of glycosylation defects on cortical pathology in CDG, we combined electrophysiological recordings using multi-electrode arrays (MEA) with comprehensive molecular profiling via multiomics, including scRNA-seq, proteomics, glycoproteomics, N-glycan imaging, lipidomics, and metabolomics. X-inactivation status was also evaluated in both iPSCs and organoids.

**Results:** ALG13-CDG hCOs revealed reduced glycosylation of proteins critical for extracellular matrix (ECM), neuronal migration, lipid metabolism, calcium ion homeostasis, and neuronal excitability. Dysregulation in related pathways was corroborated by proteomics and scRNA-seq, which also showed altered communication patterns in these pathways. Trajectory analysis revealed an inversion in neuronal development, with early inhibitory and delayed excitatory development, indicating an excitatory and inhibitory (E/I) imbalance. MEA recordings demonstrated early network hypoactivity with reduced firing rates, immature burst dynamics, and shorter axonal extensions. Despite this, transcriptomic and proteomic data revealed upregulation of excitatory receptors suggesting latent hyperexcitability. Altered lipid and sugar (GlcNAc) metabolism and skewed X-inactivation were also observed.

**Conclusions:** Our study provides the first evidence of glycosylation defects in an ALG13-CDG human cortical organoid (hCO) model and links these defects to disrupted neuronal developmental trajectories and dysregulation of key pathways essential for brain function. We identify mistimed neuronal maturation and an excitatory/inhibitory (E/I) imbalance as early drivers of network hypoactivity and immature burst dynamics, with downstream compensatory hyperexcitability that may contribute to seizure susceptibility. While specific to ALG13-CDG, these mechanisms likely extend to other glycosylation disorders with overlapping neurological features. This work offers new mechanistic insight into cortical dysfunction associated with impaired protein glycosylation and highlights potential targets for therapeutic intervention.

## Introduction

Congenital Disorders of Glycosylation (CDGs) represent a growing group of conditions affecting protein glycosylation, a process critical for protein stability and function. Among these, N-linked CDGs are caused by defects in the biosynthesis and processing of asparagine-linked glycan chains in the endoplasmic reticulum and Golgi apparatus. While CDGs often impact multiple organ systems, a substantial proportion present with predominant neurological symptoms, including developmental delay, epilepsy, and intellectual disability[1]. Asparagine-linked glycosylation 13 (ALG13) is a UDP-GlcNAc transferase that catalyzes a key step in N-linked glycosylation by adding N-acetylglucosamine (GlcNAc) to the glycan chain in the endoplasmic reticulum (**Fig 1A**). Variants in the glycosyltransferase domain of ALG13, particularly c.320A>G, p.N107S, lead to ALG13-CDG, a severe disorder of glycosylation with predominantly neurological manifestations, including developmental delay, intellectual disability, seizures, and central hypotonia [2–7]. ALG13-CDG is an X-linked condition, often with de novo mutations in females, while males experience early lethality, suggesting an X-linked dominant pattern[8].

**Figure 1:**
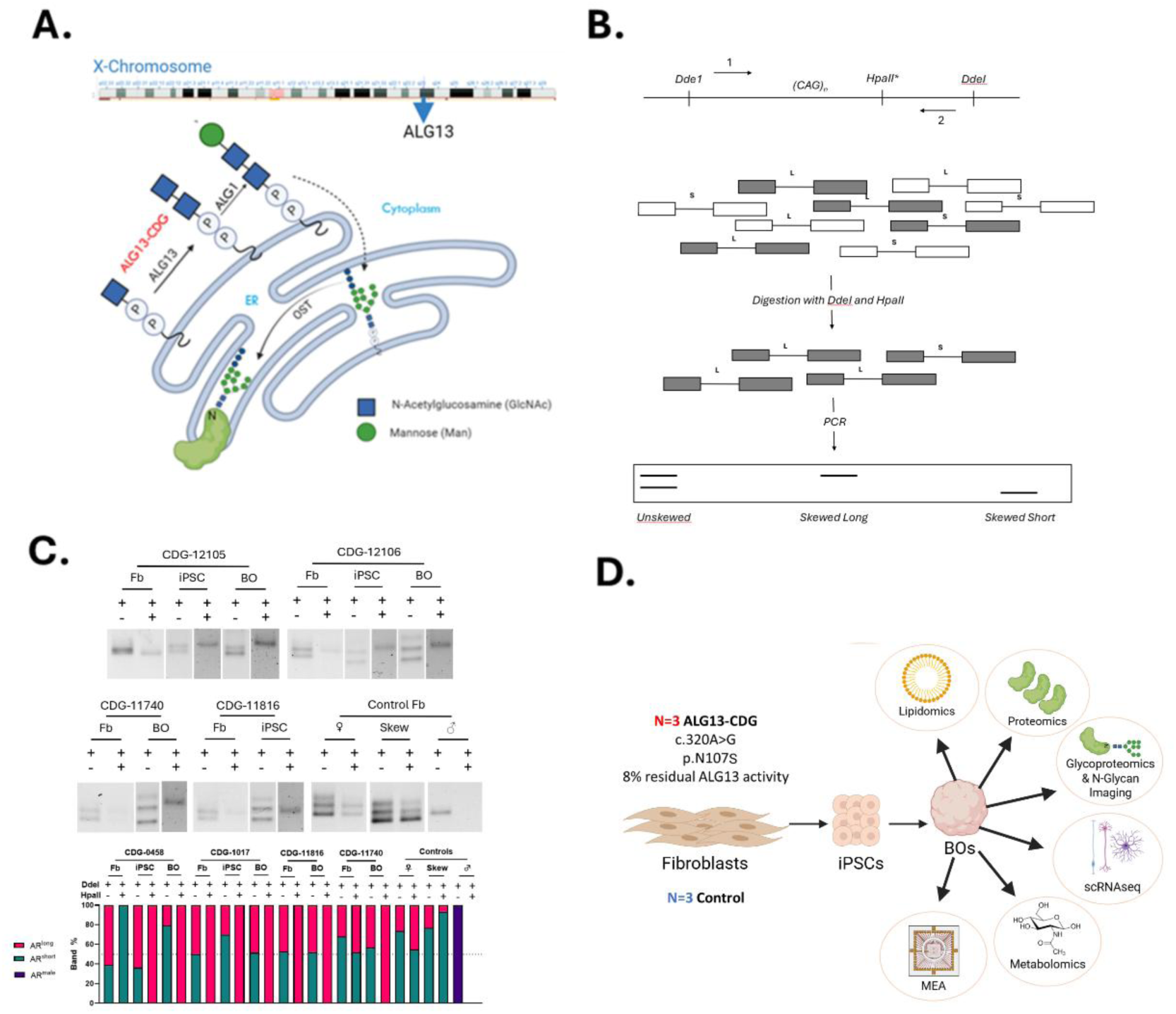
Background and Study Design. A) Protein N-linked glycosylation pathway in the ER, showing the role of ALG13. B) Schematic summary of the X-inactivation assay. Unmethylated euchromatin (white) will be digested by HpaII and will not amplify. The undigested methylated heterochromatin (grey) will be amplified, and relative abundances of the long and short polygenic AR CAG repeat regions will indicate the presence of X-inactivation skewing. C) X-inactivation skewing in ALG-13-CDG fibroblasts, induced pluripotent stem cells (iPSC) and human cortical brain organoids (hCOs). DNA agarose gel electrophoresis of PCR amplicons of androgen receptor (AR) exon1 CAG trinucleotide repeat region in gDNA incubated with DdeI and HpaII, from fibroblasts collected from ALG13-CDG fibroblasts, iPSCs and hCOs, as well as fibroblasts from a male control, and female control, and a female control with clinically assayed 90% skewed X-inactivation. The relative abundance in % of total band abundance of the polygenic alleles amplified by PCR in DNA incubated with DdeI and HpaII. F) Workflow for study.

Currently, management of ALG13-CDG symptoms is limited to seizure control with anti-epileptic drugs, ACTH treatment, and ketogenic diets, which often yield only partial relief[3–5]. Unlike other N-linked CDGs, ALG13-CDG is not detectable through standard glycosylation profiles, requiring genetic testing for diagnosis. Although studies in ALG13-deficient yeast have demonstrated glycosylation defects[8], the specific impact on brain glycosylation in humans remains unexamined. We hypothesized that ALG13-CDG may cause organ-specific glycosylation impairment, particularly within the brain, correlating with the neurological symptoms observed in affected individuals.

In this study, we reprogrammed patient-derived fibroblasts into induced pluripotent stem cells (iPSCs; **Fig 1D**) and generated three-dimensional human cortical organoids (hCOs) to study ALG13-CDG in a model system that reflects the human disease. Using this model, we performed high-resolution functional electrophysiology (multi-electrode array) and molecular analysis, including including glycoproteomics, N-glycan imaging, proteomics, scRNAseq, metabolomics, lipidomics, and, alongside assessments of X-inactivation patterns (**Fig 1B-D**).

Our findings reveal previously unidentified glycosylation defects, skewed X-inactivation, and altered gene expression and lipid profiles in ALG13-CDG, providing new insights into the disease mechanism and potential therapeutic targets.

Together, by combining molecular, transcriptomic, and functional analyses in patient-derived cortical organoids, we demonstrate that ALG13 loss-of-function leads to mistimed neuronal maturation and excitatory/inhibitory imbalance, which act as early drivers of network hypoactivity. These functional impairments likely contribute to the neurological manifestations of the disorder, including seizure susceptibility. Although this study focuses on ALG13-CDG, the developmental and circuit-level disruptions identified may represent broader, convergent mechanisms across glycosylation disorders with prominent neurological involvement. This work establishes a foundational framework for linking glycosylation-dependent molecular pathology to network-level dysfunction, offering novel ts into disease mechanisms and potential therapeutic strategies for CDGs.

## Methods

### Data Availability

The mass spectrometry proteomics data have been deposited to the ProteomeXchange Consortium via the PRIDE partner repository with the dataset identifier PXD051647. Deidentified metabolomics data have been deposited at the National Metabolomics Data Repository Metabolomics workbench and its study ID is ST003121. The scRNAseq data has been deposited to Gene Expression Omnibus (GEO) with accession number GSE266155.

### Ethics

We collected the genetic, laboratory, and clinical data of 4 patients with ALG13-CDG **(Supplementary Table 1)** enrolled in the Frontier in CDG Consortium (FCDGC) natural history study (IRB: 19–005187; NCT04199000). All patients’ skin biopsies were obtained for establishing fibroblasts as part of the standard clinical care. Informed research content was obtained from all patients included in the study. Residual samples from ALG13-CDG affected individuals (CDG-1017, CDG-0458, CDG-11740, CDG-11816) were deidentified and used for further Additional healthy fibroblasts were obtained from Coriell institute (CTRL-5400, CTRL-5381, CTRL-8399). CTRL-1363.1 and CTRL-8856 iPSCs were donated by Dr. Sergiu Pasca.

### Reprogramming of ALG13 deficient fibroblast to induced pluripotent stem cells and differentiation of ALG13 deficient Cortical organoids

The iPSCs were reprogrammed and cultured as previously described[9].ALG13-CDG iPSC’s were assessed for pluripotency markers and ability to differentiate into all 3 germ layers(**Supplementary** Figure 1). All our controls have also been previously validated for pluripotency[9]. iPSCs cells were cultured in a 10 cm plate and the organoids were developed using the dispase method as previously described[9].

### X Inactivation Skewing

Genomic DNA (gDNA) from fibroblasts, iPSCs, and cortical organoids was isolated using the QIAamp DNeasy Mini Kit (Qiagen).X-inactivation skewing was assessed using the Cutler Allen et al. method[10] with slight modifications described in **supplementary methods**.

X inactivation results in hypermethylation of one X chromosome, preventing transcription. An HpaII restriction site at c.109_112 in AR exon 1 (**Fig 1B**) allows for digestion of active, unmethylated sequences, which are not amplified by PCR. Methylated, inactivated sequences escape digestion and are amplified. By comparing the two alleles’ abundance after PCR, X-inactivation skewing can be assessed. (**Fig 1B**)

### Organoid Lysis and Protein Digestion

Five hCOs from each ALG13-CDG cell lines (CDG-11740, CDG-11816, and CDG-1017) and from control lines (CTRL-5381, CTRL-5400, and CTRL-8856.1) at Day 77 were collected and washed 3 times with DPBS, flash frozen in dry ice, and kept in −80 °C till time of assay. The samples were lysed using Bioruptor sonication device in 8M urea (in 100 mM TEABC) with 1% protease inhibitor cocktail (Thermo Scientific). Protein amount was quantified in the organoid lysates using BCA colormetric assay as per the manufacturer’s instructions (Thermo Scientific). Equal quantity of protein from both ALG13-CDG and controls were first reduced using 10 mM TCEP for 30 minutes at 55°C on a thermomixer, then alkylated with 40 mM iodoacetamide for 30 minutes in the dark at room temperature. The proteins were then digested, desalted, and tandem mass tags (TMT) labeled as previously described[9].

### Peptide fractionation, glycopeptide enrichment, liquid chromatography tandem mass spectrometry (LC-MS/MS), and data analysis

Twenty percent of the peptides were fractionated using bRPLC for proteomics and 80% of the peptides were fractionated using size-exclusion chromatography (SEC) for glycoproteomics as previously described[9]. LC-MS/MS for glycoproteomics and proteomics was conducted as previously described[9] with slight alterations provided in **supplementary methods**. Data analysis for glycoproteomics and proteomics was performed as described previously[9].

### Tissue Preparation,processing, and anlaysis for MALDI-MSI

CDG-11740, CDG-11816, CDG-1017, CTRL-1363.1, CTRL-8399, and CTRL-8856.3 hCOs at day 145 were preserved as described[11]. Spheroids were sectioned at 12um onto glass slides. Samples were prepared similarly to a standardized protocol as described[12]. Tissue samples were analyzed on a timsTOFfleX MALDI-QTOF as described in **supplementary method**.

### Single Cell Suspension Formation, Library Preparation, and Sequencing for scRNAseq

At day 92, four organoids per cell line (CDG-0458, CDG-1017, CDG-11740, CTRL-8856.3, CTRL-1363.1, CTRL-8399) were dissociated into single cells using the Miltenyi Biotec Neurosphere Dissociation Kit, with a modified 20-25 minute incubation at 37°C. Cells were resuspended in 1x PBS + 0.04% BSA and sent to the Genome Analysis Core (GAC) for Single Cell partitioning. Library preparation and sequencing details are in **supplementary methods**.

### scRNAseq Data Analysis

Details on how fastq files were used to generate an integrated Seurat object can be found in **supplementary methods**.After integration, data was scaled with ScaleData, and PCA was performed. Clustering analysis using FindNeighbors, FindClusters, and UMAP (dimensions 1:20) identified eight distinct clusters. Cluster identities were determined using FindAllMarkers and FindConservedMarkers and matched with literature[13–16]. 7 of the clusters were identified according to their markers **(SFig 4)** One cluster remained unclassified due to lack of definitive markers.

After cluster identification, differential gene expression analysis was performed by aggregating data into pseudobulk samples per organoid line and running DESeq2 with the apeglm method[17] to identify significant transcript changes using padjusted values <0.05. The integrated Seurat object was split into ALG13-CDG and control sample objects for trajectory analysis using Monocle3[18]. Following this analysis, the previously unidentified cluster was reclassified as multipotent progenitor cells (MPC) due to its branching toward inhibitory and astrocyte progenitor cell populations.

#### CellChat Analysis

The cell–cell interactions between the different cell types in the organoid were evaluated using CellChat (R package)[19]. First, the split Seurat object was used to generate CellChat objects for Controls and ALG13-CDG. The CellChatDB.human, which is a database that contains information about known interactions between receptors, ligands, and cofactors, was integrated into the CellChat objects to allow the determination of cell-cell communication networks. The CellChat data was processed as previously described[19].

### Organoid Preparation and Metabolite measurement for Metabolomics

5 cortical organoids from each ALG13-CDG cell lines (CDG-11740, CDG-11816, and CDG-1017) and from control lines (CTRL-5381, CTRL-5400, and CTRL-8856.1) were collected and washed 3 times with DPBS, flash frozen in dry ice, and kept in −80°C till time of assay. At time of assay, hCOs were washed with saline and weighed. 350 µL of ice-cold extraction buffer (80 % MeOH, Internal Standards) and a few sonication beads were added to the sample and placed in Bioruptor sonication device for lysing. After lysing, the metabolite extraction, LC-MS, and normalization were conducted as previously described[9].

### Lipid extraction from organoids

Untargeted lipidomics analysis was performed on cortical organoids from three ALG13-CDG lines (CDG-11740, CDG-12105, CDG-12106) and five control lines (CTRL-5400, CTRL-5381, CTRL-8399, CTRL-1363.1, CTRL-8856). Deuterated lipids were added prior to conducting a modified Bligh and Dyer lipid extraction[20]. The dried organic layer was reconstituted in chloroform (1:3 v/v) for LC-MS/MS analysis.

### LC-MS/MS analysis of lipids

Untargeted lipidomics analysis was performed on an Orbitrap IQ-X Tribrid mass spectrometer (Thermo Fisher Scientific) connected to Vanquish Horizon UHPLC (Thermo Fisher Scientific). AcquireX Deep Scan, a data-driven MS/MS acquisition approach was used to maximize the coverage of lipids as previously described^. The^ lipids were separated as previously described. Full scan MS (m/z 250–1500 in positive ion mode and 250-1600 m/z in negative ion mode) was acquired at a resolution of 120,000 (at m/z 200) while MS/MS scans were acquired at a resolution of 15,000 (at m/z 200).

### Data Analysis for Lipidomics

Lipids were identified and peak areas were integrated using LipidSearch 5.1 (Thermo Fisher Scientific) based on precursor m/z and MS/MS or MS^3^ spectra. Peak areas were normalized to deuterated internal standards added before extraction and to total lipid levels. Statistical comparisons between ALG13-CDG and controls were conducted using Student’s t-test.

### Plating of cBOs on Multi-Electrode Array (MEA) and activity, network, and axon recording

Electrophysiological recordings were conducted using the 6-well high-density MEA MaxTwo system (HD-MEA) from MaxWell Biosystems. Wells were prepared following the MaxWell “Brain Organoid Plating Protocol, V2.0,” with minor modifications. Briefly, 6-well MEA plates were incubated in 1% Tergazyme solution for 2 hours at room temperature (RT), washed three times with distilled water, sterilized in ethanol for 30 minutes, and washed again three times with distilled water. Each well received 1 mL of Neurobasal-A (NBA) medium and was incubated for 48 hours for pre-conditioning.

After incubation, 50 µL of 0.07% poly(ethyleneimine) (PEI) was added to each well and incubated for 1 hour. Wells were then washed three times with distilled water and air-dried for 1 hour at RT. Next, 50 µL of 0.04 mg/mL laminin in NBA was added and incubated overnight. Laminin was aspirated, and cortical brain organoids were placed directly on the electrodes without media for 2–5 minutes. Subsequently, 50 µL of 0.04 mg/mL laminin in NBA was slowly added to each well, and the plates were incubated for 2 hours. For each organoid line, four MEA wells were prepared. After the 2-hour incubation, 1 mL of NBA was gently added to each well. The following day, half of the media was replaced with fresh NBA.

Organoids were maintained by changing the media every 3–4 days and at least 24 hours before MEA recordings on Day 120,124,133, and 140. Recordings were performed using MaxLab software. The built-in “Activity Scan” protocol was used to identify active areas and assess firing rates. All parameters were kept at default except the recording duration, which was extended to 60 seconds to better detect slower activity. This was followed by the “Network Assay” protocol to evaluate neuronal network firing, using the neuronal units parameter. Lastly, the “Axon Tracking Assay” was performed using block parameters to detect axonal signal propagation.

For each protocol, wherever feasible data were averaged across the four wells per sample. Statistical significance was determined using Two-Way Anova and Šídák post hoc tests.

## Results

### Study Cohort

The study included four ALG13-CDG patients carrying the common de novo c.320A>G, N107S variant. Two patients (CDG-0458 and CDG-10175) were previously reported, while two (CDG-11740 and CDG-11816) are newly described. All patients presented with neurological symptoms, including developmental delay, seizures (infantile spasms), intellectual disability, and delayed or absent speech (**STable 1**). CDG-11740 achieved seizure control on a ketogenic diet, but others did not. Additionally, CDG-1017 and CDG-11816 exhibited central hypotonia, and CDG-11816 and CDG-11740 had ophthalmological abnormalities. CDG-11740 also had hearing loss (**STable 1**).

### X-Inactivation Skewing in ALG13-CDG

Contrary to previous reports suggesting random X-inactivation, three of four patient fibroblast lines showed complete X-inactivation skewing, with CDG-11740 as the exception, exhibiting random inactivation (48% and 52% AR allele methylation;**Fig 1-B-C**). However, cortical organoids derived from all ALG13-CDG iPSCs displayed complete skewing. Control lines confirmed the reliability of this methodology.

### Proteomic Analysis of ALG13-CDG hCOs

Global proteomic analysis of three ALG13-CDG and three control cortical brain organoid lines identified 8,750 proteins, with 321 significantly altered (**Fig 2A-B**). Altered proteins included those critical for neuronal function, ER stress, oxidative stress, ECM integrity, and neuronal migration (**Fig 2A-F**). Specifically, SELENOS, TXLNG, PSMD10, and APIP were dysregulated, impacting ER stress-related apoptosis, while proteins such as OTX2, LGALS3, RDH10, and VGF, essential for brain development, also showed changes. Dysregulation of SELENOS, SELENBP1, and VGF indicated altered lipid metabolism, and downregulation of ITPRIP and PDGFRB indicated calcium ion homeostasis dysregulation.

**Figure 2.**
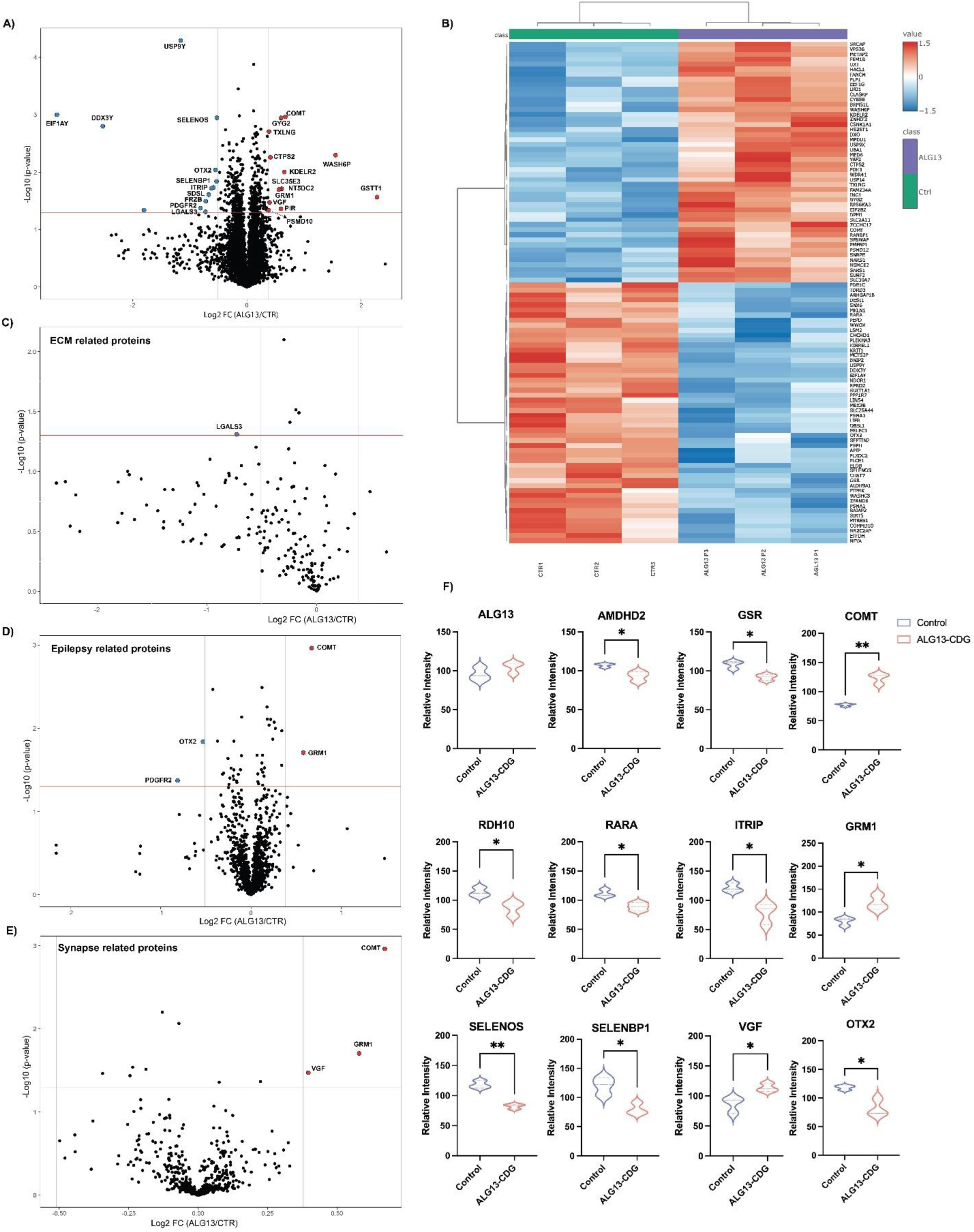
Protein changes in ALG13 deficient hCOs. **A)** Volcano plot depicting the differentially expressed proteins in ALG13-CDG. X-axis is log_2_ fold-change (ALG13-CDG/controls) and Y-axis is the negative logarithm of p-value from a t test for significance as indicated. The horizontal dashed red line represents the cutoff for significance (<0.05). Proteins names are provided for peptides that have p value < 0.05 and fold change of 1.3. Left-shifted proteins are downregulated (blue dots) and right shifted proteins are upregulated (red dots) in ALG13 deficient brain organoids. **B)** Heatmap of significantly altered proteins with p value < 0.05. Volcano plots depicting the differential expression of proteins related to (C) extracellular matrix (ECM), (D) epilepsy, (E) synapse. **F)** Violin Plots for specific proteins (p value<0.05).

In examining ECM-associated dysregulation, proteins like LGALS3, HS3ST1, and KRIT1, vital for neuronal development[21–23], were downregulated (**Fig 2A, 2B, 2F**). Additionally, RARA and RDH10, important neuronal migration, were downregulated. Epilepsy-related proteins, including upregulation of GRM1 and COMT and downregulation of PDGFRB and OTX2, were noted (**Fig 2D**). Synaptic function proteins, such as GRM1, COMT, and VGF, showed upregulation, and AMDHD2, crucial for GlcNAc catabolism[24], was downregulated.

Notably, Y-linked genes, like EIF1AY, appeared downregulated in ALG13-CDG hCOs, consistent with the sex distribution in our cohort and known Y-linked gene downregulation in females[25]. Differences between male and female controls were negligible, suggesting no gender-related bias in our findings.

### ALG13-CDG hCOs Exhibit Distinct N-Glycosylation Remodeling

Our N-glycoproteomics analysis of ALG13-CDG hCOs identified 2,890 glycopeptides with 282 unique N-glycan structures across 510 sites on 412 glycoproteins.

Approximately 25% of these glycans were high mannose, with Man5 and Man8 as the most common types, while the remaining 75% were complex/hybrid (**Fig 3A, SFig 5A**). Among 66 glycopeptides from 45 glycoproteins showing significant glycosylation changes, 60 glycopeptides exhibited reduced abundance in ALG13-deficient fibroblasts, indicating a broad glycosylation deficit across patients (**Fig 3B, SFig 5B**). Partial Least Squares Discriminant Analysis (PLS-DA) further confirmed glycopeptide-level distinctions between ALG13-CDG and control hCOs (**SFig 5C**).

**Figure 3.**
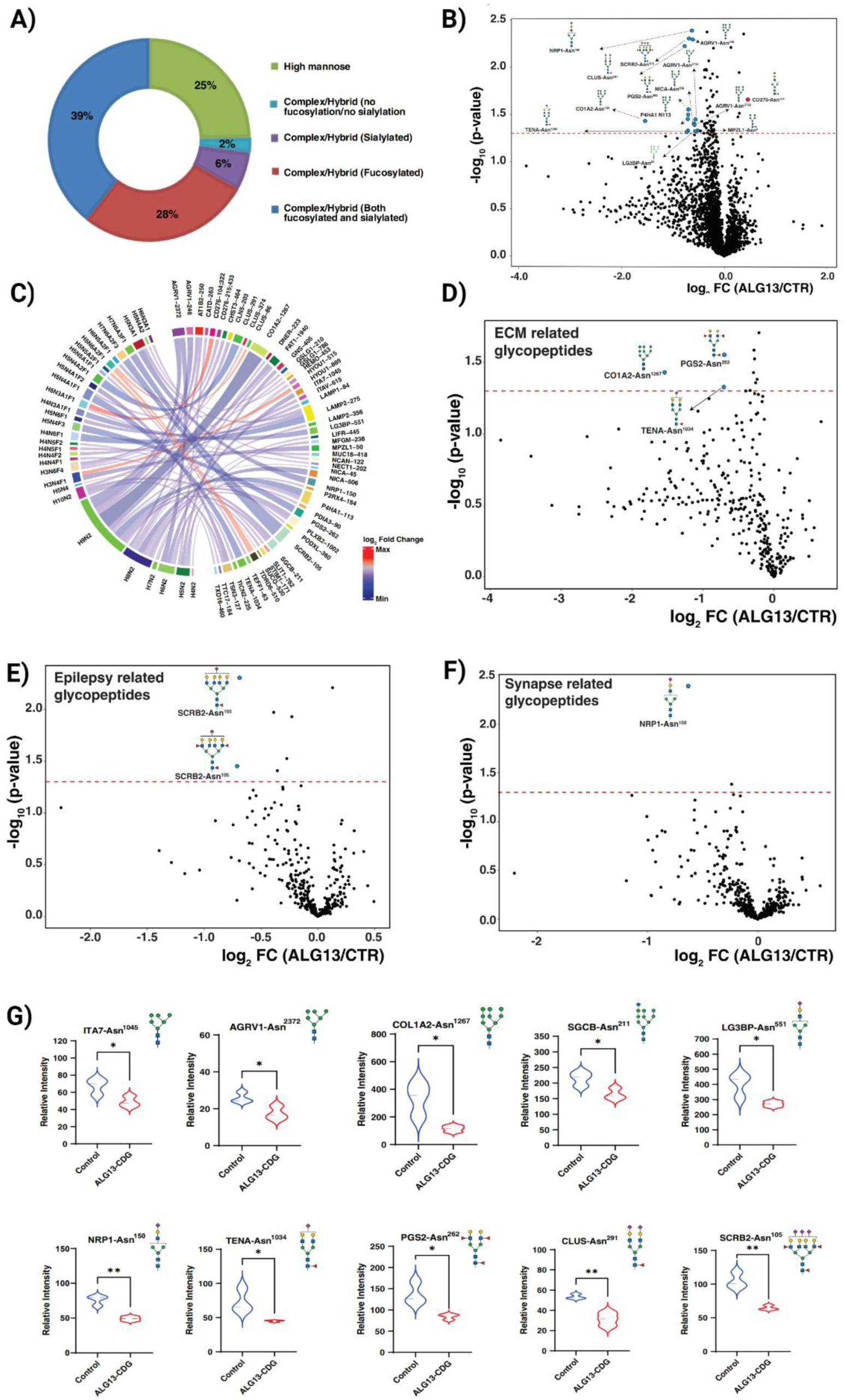
Site-specific glycosylation changes and glycosylation remodeling of different pathways in patient-derived ALG13-CDG hCOs. **(A)** All quantified glycopeptides were categorized based on their glycans, with corresponding percentages provided. **(B)** Volcano plot depicting the differentially expressed glycopeptides in ALG13-CDG. X-axis is log_2_ fold-change (ALG13-CDG/controls) and Y-axis is the negative logarithm of p-value from a t test for significance as indicated. The horizontal dashed red line represents the cutoff for significance (<0.05). Some of the changing glycopeptides are marked in red circles and glycoproteins’ names, glycosylation sites and glycan structures are drawn. **(C)** Differential chord diagram depicting significantly changing glycopeptides with the proteins and site of attachment in ALG13-CDG as compared to control hCOs. Proteins with different glycosylation sites are indexed on the right of the diagram and connected via chords to respective identified glycan structures on the left. Asn^x^ represents the asparagine at amino acid site “x” in the corresponding protein sequence. The fold-change pattern is color coded. Putative structures are shown using Symbol Nomenclature for Glycans (SNFG). p < 0.05 (*), p < 0.01 (**).Volcano plots depicting the differential expression of glycopeptides from proteins related to (D) extracellular matrix (ECM)_(E) epilepsy (F) synapse. X-axis is log2 fold-change (ALG13-CDG/controls) and Y-axis is the negative logarithm of p-value from a t test for significance as indicated. The horizontal dashed red line represents the cutoff for significance (<0.05). Some of the changing glycopeptides are marked in red circles and glycoproteins’ names, glycosylation sites and glycan structures are drawn. (G) Violin plots showing relative intensities of several downregulated regulated glycopeptides in ALG13-CDG derived from numerous proteins associated with these pathways. p < 0.05 (*), p < 0.01 (**).

**Figure 4.**
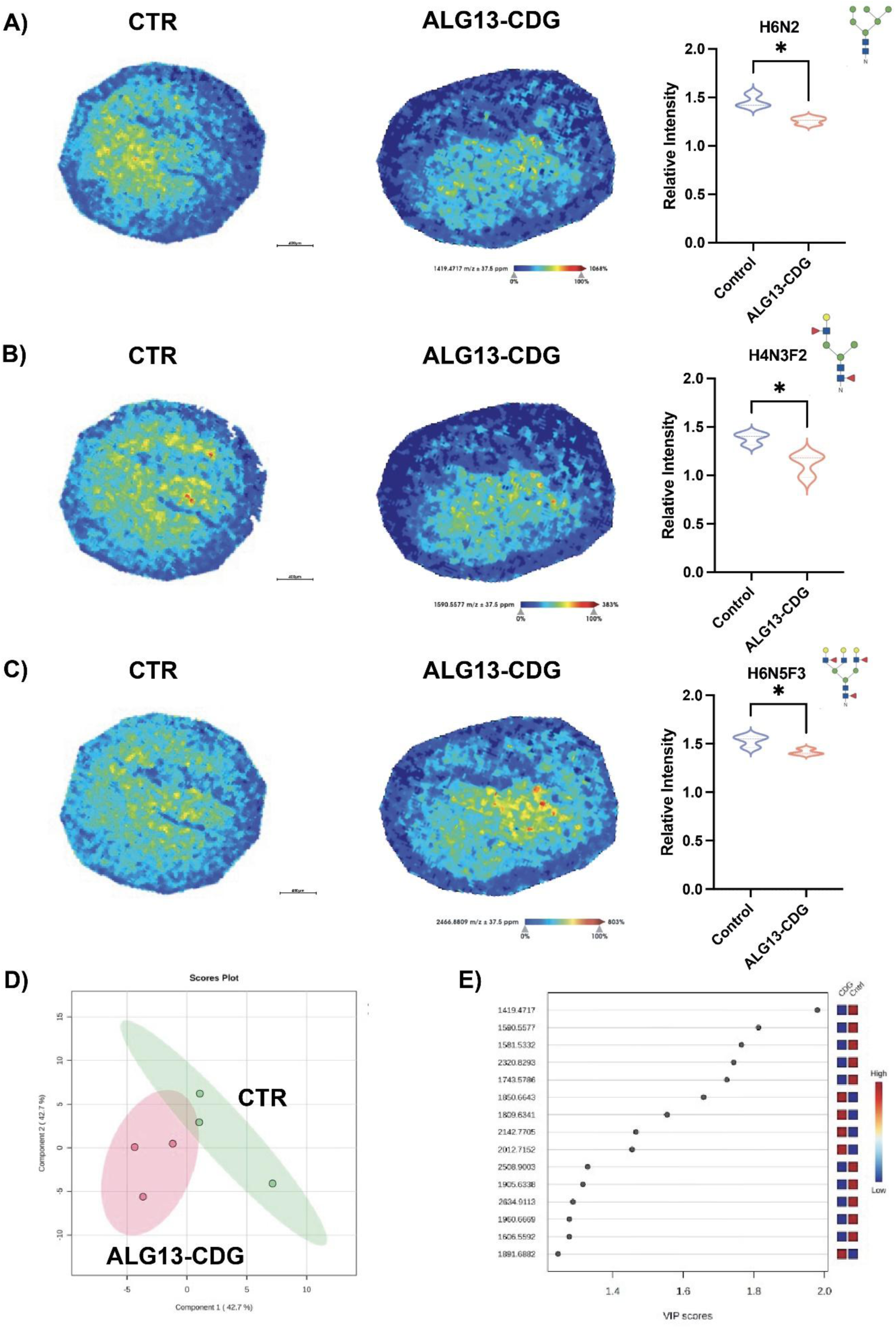
N-glycomic comparison of ALG13 deficient hCOs to controls. MALD-MSI representative images of m/z **A**)1419.4717 H6N2, **B**)1590.5577 H4N3F2, and **C**) 2466.8809 H6N5F3 N-glycans for control and ALG13 deficient cortical organoids with corresponding relative abundance and N-glycan structure on the right. **D**) Partial least squares discriminant analysis of the top 15 N-glycans separating the groups with corresponding **E**) VIP Scores plots. Shaded areas are 95% confidence intervals. Each point represents 1 patient. Intensity gradient from blue (least abundant) to red most abundant. Scale bars are below images. *p<0.05, using students T-tests.

**Figure 5.**
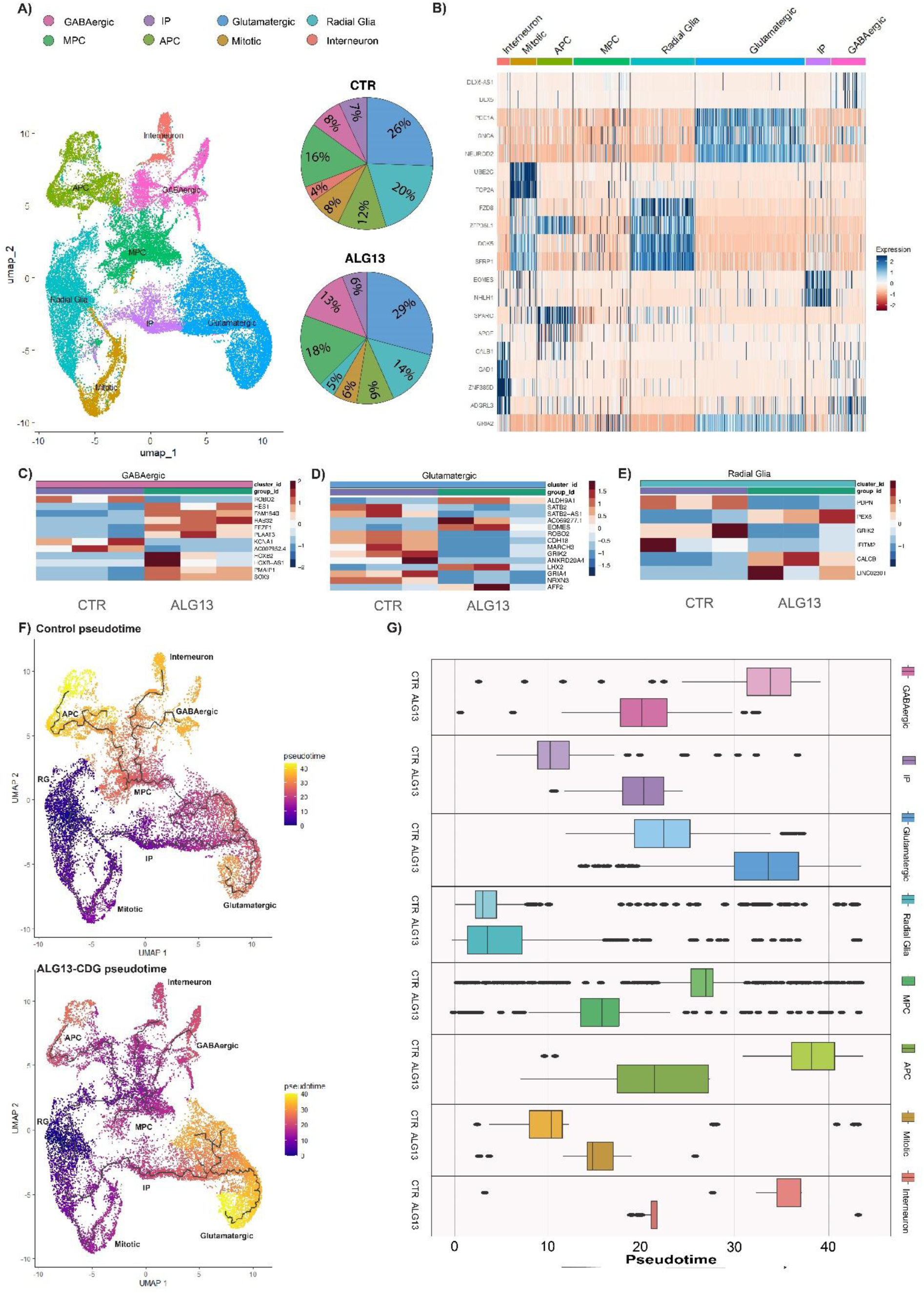
scRNAseq comparison between ALG13-CDG and Controls show an irregular developmental trajectory and alteration in transcripts important for brain development and function. **A)** UMAP of integrated scRNAseq data that shows the different cell clusters identified in both ALG13-CDG and controls. Percent cell in each cluster for ALG13-CDG and Control hCOs. **B)** Heatmaps of top markers used to identify cluster. **C-E)** Heatmaps of differential gene expression analysis for clusters with significant gene changes. **F)** Trajectory analysis graph for control and ALG13-CDG hCOs, showing the development trajectory of different cell types through color coding associated with pseudotime. **G)** Boxplots quantifying the pseudotime per cluster in control and ALG13-CDG hCOs.

A subset of 23 hypoglycosylated glycopeptides was associated with pathways essential for lysosomal function, lipid metabolism, neuronal migration, apoptosis, and synaptic function[8, 26–32] (**Fig 3B**). Although not all glycopeptides reached statistical significance, proteins involved in ECM, epilepsy, and synapse-related pathways exhibited reduced glycosylation, potentially contributing to neuronal and synaptic instability (**Fig 3D–G**). Given that GlcNAc-bisected glycans are predominantly brain-specific, we noted approximately 21% (609 glycopeptides) were GlcNAc-bisected, with a global reduction in ALG13-CDG hCOs, prominently affecting MUC18, PODXL, and TEF1 (**SFig 5D**).

### N-glycomic Analysis Reveals Distinct Differences in ALG13-CDG hCOs

Glycan imaging identified 57 N-glycans in both ALG13 deficient and controls hCOs **(Fig S2A)**. Further univariate partial least squares discriminant analysis revealed N-glycan species of high mannose, bisecting, and multiantennary N-glycans were prominent discriminators between ALG13-CDG and control (**Fig 4A-B**). Notably, brain-specific N-glycan species[33] m/z 1257.4173 H5N2, m/z 1688.6083 H3N5F1, and m/z 1996.7276 H4N5F2 were present in both groups without significant differences **(Fig S2B-D)**, supporting the validity of our hCO model. **(Fig S2B-D)**. Notable decreases were observed in m/z 1419.4717 H6N2, m/z 1590.5577 H4N3F2, and m/z 2466.8809 H6N5F3 in ALG13-CDG hCOs compared to controls (**Fig 4A-C**). These differences align with glycoproteomics data indicating altered H6N2 glycosylation in proteins linked to neuronal function, metabolism, and migration.

### scRNAseq analysis reveals irregular developmental trajectory, cell communication networks, and alteration in transcripts associated with brain development and function in ALG13-CDG hCOs

To assess gene expression differences in ALG13-CDG hCOs, we conducted single-cell RNA sequencing (scRNA-seq) and identified eight cell types: Radial Glia (RG), Mitotic Radial Glia, Intermediate Progenitors (IP), Glutamatergic Neurons, GABAergic Neurons, Interneurons, Multipotent Progenitor Cells (MPCs), and Astrocyte Progenitor Cells (APCs) (**Fig 5A, SFig 4**). Cluster analysis revealed no significant differences in cell percentages between ALG13-CDG and control hCOs (**Fig 5A**). However, differential gene expression analysis showed significant dysregulation in RG, Glutamatergic, and GABAergic cells, particularly in genes linked to cell migration, apoptosis, axon guidance, oxidative stress, lipid metabolism, and calcium ion regulation (**Fig 5C-E**). In neurogenesis, HES1 and SOX3 were downregulated in the GABAergic cluster, and EOMES and LHX in the Glutamatergic cluster (**Fig 5C-D**). PLAAT3 and PEX6, essential for lipid metabolism, were downregulated in the GABAergic and RG clusters, respectively (**Fig 5C, 5E**). Additionally, the long non-coding RNA AC007952.4, associated with oxidative stress, was upregulated in ALG13-CDG, alongside altered transcripts linked to neuronal migration (*ROBO2, SATB2, FEZF1, PDPN*) and upregulation of KCNA1 and glutamate receptors (*GRIK2, GRIA4*) (**Fig 5C-E**). CellChat analysis indicated dysregulation in cell communication networks, with heightened activity in collagen, cadherin, and EGF pathways, which may impact ECM function, neuronal migration, and calcium ion homeostasis (**SFig 6, Fig 6A-C**). Trajectory analysis showed delayed development of the glutamatergic population in ALG13-CDG hCOs, while MPC, APC, and inhibitory neuron populations developed earlier than in controls (**Fig 5F-G**).

**Figure 6:**
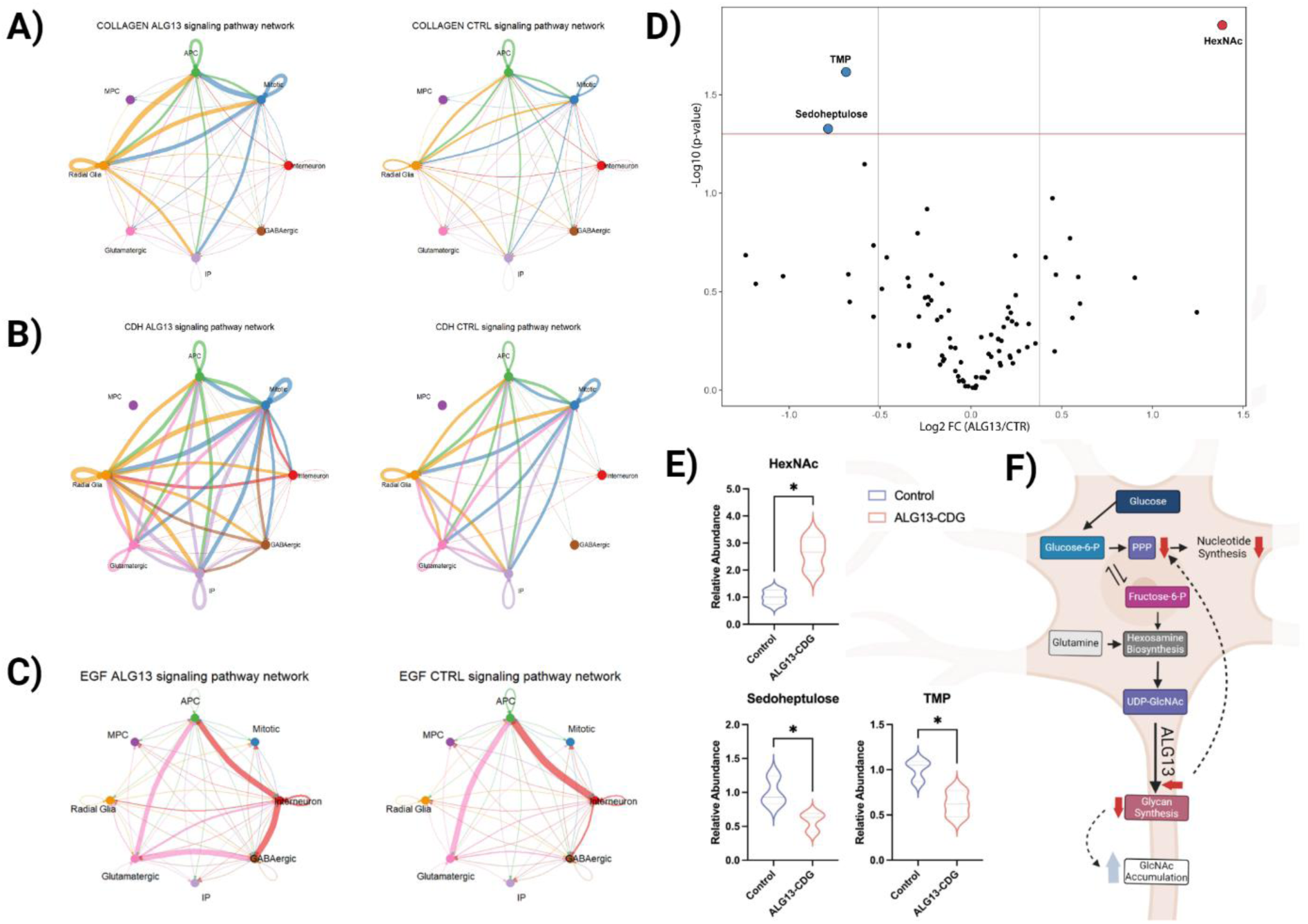
Remodeling of cell communication networks and metabolites in ALG13 deficient hCOs. A-C) Chord Diagrams for specific signaling pathways crucial for extracellular matrix function(A), neuronal migration (B), and calcium ion homeostasis (C). Chords connecting the cell nodes indicate communication between the cell types in regard to the pathways listed. The width of the cord correlates with the weight/amount of communication. **D)**Volcano Plot comparing ALG13 deficient brain metabolites to controls. X-axis is log2 fold-change (ALG13-CDG/controls) and Y-axis is the negative logarithm of p-value from a t test for significance as indicated. The horizontal dashed red line represents the cutoff for significance (<0.05). Metabolites names are provided for metabolites that have p value < 0.05 and fold change of 1.3. **E)** Violin plots for significantly changing metabolites (p<0.05). **F)**Schematic representation of the metabolic alterations in ALG13 deficient hCOs.

### Metabolic Analysis Reveals Limited Alterations in ALG13-CDG hCOs

Congenital disorders of glycosylation (CDGs) often induce global metabolic rewiring[9, 34]. To assess ALG13 deficiency’s impact on metabolism, we performed targeted metabolomics, focusing on GlcNAc-related pathways, including glycolysis, hexosamine biosynthesis, glutamine metabolism, and the pentose phosphate pathway (PPP). Although many metabolites were quantified, ALG13-deficient hCOs showed minimal metabolic differences from control (CTR) hCOs (**Fig 6D, SFig 7**). Notably, PPP metabolites like TMP and D-Sedoheptulose were downregulated, while HexNAc levels (GlcNAc, GalNAc, and ManNAc) were significantly elevated (**Fig 6E**).

**Figure 7.**
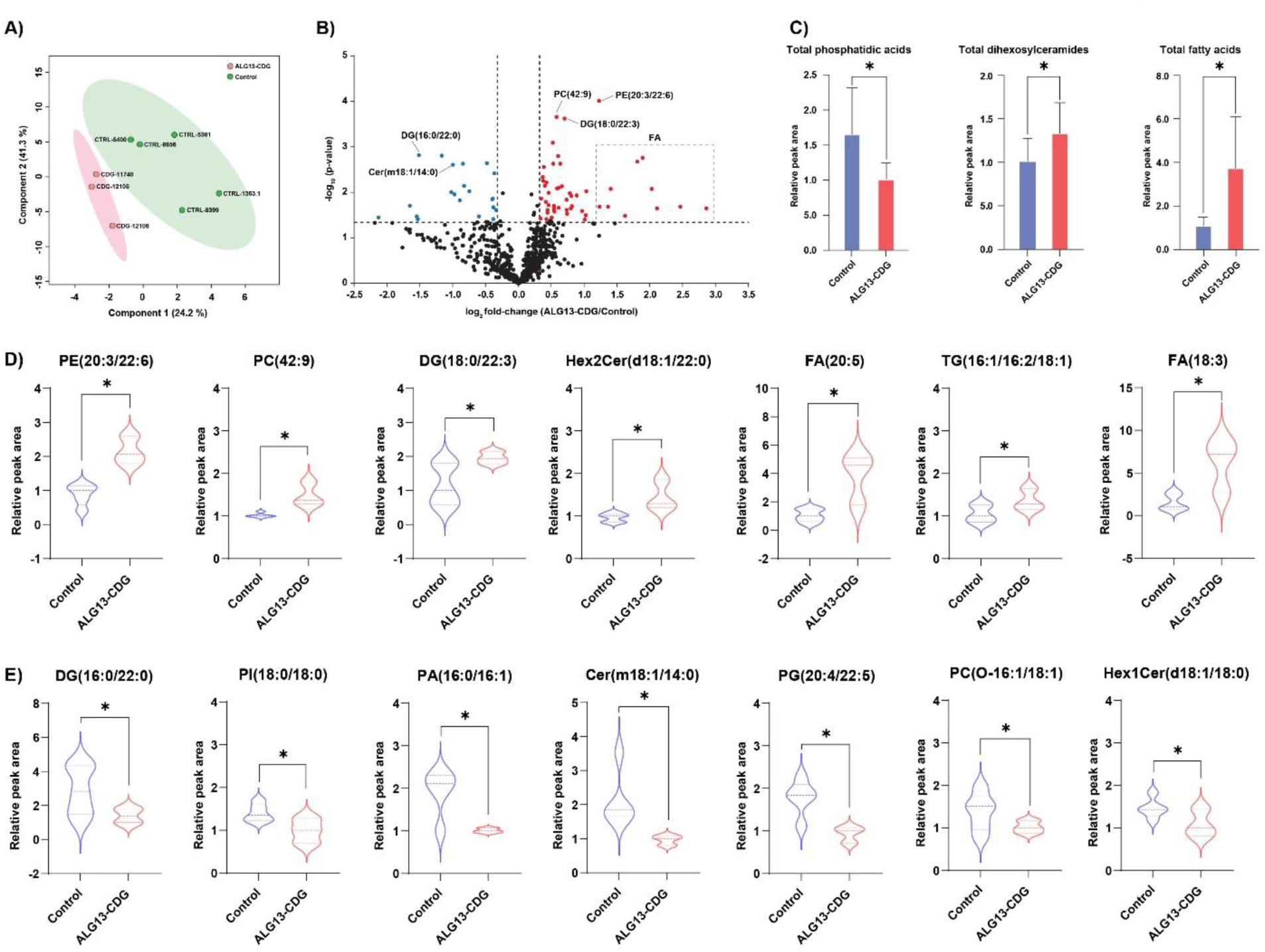
Lipid alterations in ALG13-CDG hCOs. **A)** Scores plot from partial least squares discriminant analysis of all identified lipids. **B)** Volcano plot depicting lipidomics changes in ALG13-CDG hCOs. PE(20:3/22:6), PC(42:9), DG(18:0.22:3) and several FA species were significantly increased, while DG(16:0/22:0) and Cer(m18:1/14:0) were decreased significantly. C) Total levels of subclasses of lipids with significant alterations in ALG13-CDG hCOs. **D)** Violin plots showing lipid species which are significantly increased in ALG13 deficient hCOs. **E)** Violin plots showing the lipids which are significantly decreased in ALG13 deficient hCOs. Abbreviations: PE, phosphatidylethanolamine; PC, phosphatidylcholine; DG, diglyceride; Hex2Cer, dihexosylceramide; FA, fatty acid; TG, triglyceride; PI, phosphatidylinositol; PA, phosphatidic acid; Cer, ceramide; PG, phosphatidylglycerol; Hex1Cer, monohexosylceramide.

### Global lipidomic analysis reveals lipid alteration in ALG13-CDG hCOs

Global lipidomic profiling of ALG13-CDG and control hCOs quantified 676 lipids across 23 phospholipid subclasses.. Total lipid levels for each subclass and individual lipid species abundances are provided in **Supplementary Tables 2 and 3**. Distinct lipid profiles separated ALG13-deficient hCOs from controls (**Fig 7A**), with significant lipid alterations highlighted in a volcano plot (**Fig 7B**). In ALG13-CDG hCOs, levels of lipids such as PE(20:3/22:6), PC(42:9), Hex2Cer(d18:1/22:0), and several FA species were increased, while DG(16:0/22:0) and Cer(m18:1/14:0) were decreased. Total PA levels significantly decreased, whereas Hex2Cer and FA increased in ALG13-deficient hCOs (Fig 7C). Overall, 60 lipid species, including PE, Hex2Cer, and FA, showed significant increases (fold-change >1.25, p < 0.05) in ALG13-CDG (**Fig 7D**), while 27 species decreased (fold-change <0.8, p < 0.05), as shown in Fig 7E.

### ALG13-CDG Cortical Organoids Exhibit Reduced Activity and Impaired Neuronal Network Maturation and Activity

ALG13-CDG hCOs showed significantly reduced 90th percentile firing rates (**Fig 8B**,Group Effect p = 0.0158), indicating that even the most active neurons in ALG13-CDG fire less than controls and are hypoactive. Spike per burst pr std (**Fig 8C**, Group Effect p = 0.0158) is also diminished in ALG13-CDG, which indicates a more immature network as the variability in spike numbers within bursts is uniform and weaker. The burst peak firing rate 10^th^ percentile dynamics is also abnormal (group × time interaction, p = 0.0080), indicating hypoactivity over time and worsening network coordination.

**Figure 8.**
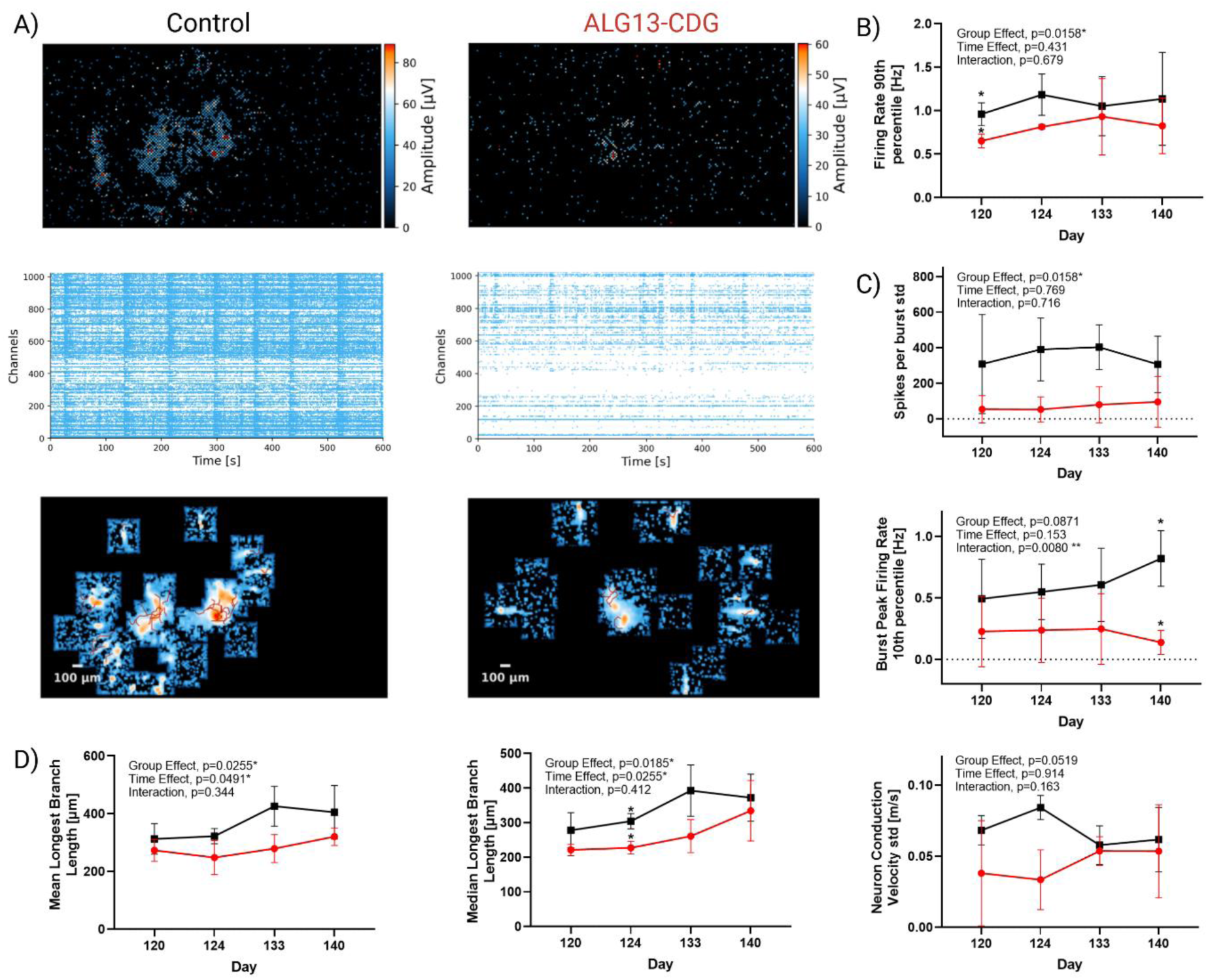
Neuronal network deficits in ALG13-CDG hCOs. A) Representive amplitude maps, raster plots, and axon tracking maps from Day 120-140 hCOs for control (left) and ALG13-CDG (right). Line graphs summary of MEA data from Day 120,124,133, and 140. B)Summary for activity metric Firing Rate 90^th^ Percentile [Hz].C) Summary for network metrics Spike per Burst per std and Burst Peak Firing Rate 10^th^ percentile [Hz]. D) Summary for axon metrics Mean Longest Branch Length [um], Median Longest Branch Length [um], and Neuron Conduction Velocity [m/s]. Data were analyzed by two-way ANOVA with Šídák post hoc correction. *p < 0.05; **p < 0.01. Error bars indicate SEM.

Axonal signal tracking showed shorter mean and median longest branch lengths (**Fig 8D**) in ALG13-CDG (Group Effect p = 0.0255 and p = 0.0185, respectively), indicating neuronal immaturity and deficits in axon extension. A trend toward decreased conduction velocity variance (Group Effect p = 0.0519) was also seen in ALG13-CDG (**Fig 8D**), which is consistent with immature neurite outgrowth and signal propagation.

## Discussion

This study presents the first biochemical evidence of brain-specific N-glycosylation abnormalities in ALG13-CDG, addressing a longstanding gap in understanding the molecular basis of this disorder’s neurological symptoms. While individuals with ALG13-CDG often have normal serum glycosylation profiles, our findings reveal significant glycosylation defects in brain tissue, aligning with the predominantly neurological phenotype. Through a comprehensive multi-omics profiling, we have identified key pathways potentially contributing to seizures, intellectual disability, and developmental delays in ALG13-CDG, paving the way for future research and treatment strategies.

### X-Inactivation Skewing in ALG13-CDG

Our study identified significant X-inactivation skewing in three of four ALG13-CDG fibroblast lines, contrasting with previous reports of random X-inactivation in these patients[35]. Extensive X-inactivation skewing was also observed in iPSC-derived cortical organoids from ALG13-CDG patients, highlighting the potential role of X-linked epigenetic regulation in disease pathology. The consistency of this inactivation across cell types supports the robustness of our findings and provides insight into how X-inactivation dynamics may modulate gene expression and impact the severity of neurological symptoms in ALG13-CDG.

### Extracellular Matrix Dysregulation, Neuronal Migration, and axon extension dysregulation

Our results demonstrate that hypoglycosylation of key ECM components, including collagen (CO1A2) and proteoglycan (PGS2, **Fig 3**), could disrupt ECM integrity, an essential factor for neuronal migration and axon extension during cortical development. Neuronal migration depends on interactions between ECM proteins and radial glial cells, which help direct neurons to their proper locations[28]. Altered glycosylation of integrins (ITA7, LG3BP) and the upregulation of PDPN may further destabilize these critical cell-ECM interactions[29], potentially explaining the developmental delays[27] and seizure phenotypes[36] observed in ALG13-CDG. Additionally, reduced levels of ECM-related proteins, such as RDH10 and RARA(**Fig 2B,2F**), linked to microcephaly, indicate that ECM dysregulation may contribute to the structural brain abnormalities reported in ALG13-CDG[37]. Furthermore, as the ECM is crucial for axon extension, ECM disruption may also underlie the reduced axonal extension in ALG13-CDG (**Fig 8A, 8D**)

### ER and Oxidative Stress

We observed that several ER stress-related genes, including TXLNG, PSMD10, and SELENOS (**Fig 2A**), are dysregulated in ALG13-deficient hCOs. This suggests that ALG13-CDG may contribute to protein misfolding and the unfolded protein response, which can exacerbate oxidative stress due to reactive oxygen species production[38]. Given the brain’s high vulnerability to oxidative stress[39, 40], these findings imply that ER stress and oxidative imbalance may underlie the seizures, developmental delays, and cognitive impairments associated with ALG13-CDG. Our scRNAseq data, showing upregulation of oxidative stress markers (**Fig 5C**) in GABAergic clusters, reinforces this theory and highlights potential targets for therapeutic intervention to manage oxidative stress in these patients.

### Calcium Ion Homeostasis

Our analysis identified hypoglycosylation of SGCB and AGRV1, proteins important for maintaining calcium homeostasis[30, 31], in ALG13-CDG. Disrupted calcium regulation has significant implications for neuron function, as calcium signaling is crucial for cognitive processes[41], synaptic plasticity, and neuronal excitability. Further supporting this, our proteomics data revealed downregulation of ITRRIP and PDGFRB(**Fig 2A**), both of which play roles in calcium signaling and have been associated with epilepsy[41]. The disruption of calcium signaling may, therefore, contribute to the complex neurological symptoms in ALG13-CDG, including seizures, developmental delays, and intellectual disability. Furthermore, altered EGF signaling between glutamatergic and GABAergic clusters (**Fig 6C**) in ALG13-CDG suggests a broader impact on calcium-dependent developmental pathways[42], potentially affecting excitatory and inhibitory neuron development (**Fig 5C-G**).

### Lipid Metabolism Dysregulation

Our lipidomic profiling identified significant disruptions in lipid metabolism in ALG13-CDG. Notable increases in lipid species(**Fig 7B-D**), including FA(22:1), FA(22:4), and FA(22:6), suggest peroxisomal dysfunction[43], supported by decreased PEX6 transcript levels(**Fig 5C**). Elevated lactosylceramide (Hex2Cer) levels, linked to oxidative stress and mitochondrial dysfunction[44], may contribute to the seizure and developmental delay phenotypes observed in ALG13-CDG. Decreased levels of PA, essential for dendritic spine maturation and synaptic plasticity[45], further suggest a connection between lipid metabolism dysregulation and the intellectual disability phenotype.

Proteomics data also indicated downregulation of AMDHD2 (**Fig 2F**), implicating disrupted GlcNAc catabolism and an increase in intracellular GlcNAc levels(Fig 6), both of which are associated with lipid dysregulation and may have additional impacts on neuronal function.

### Neuronal Hyperexcitability and Seizure Phenotypes

Neuronal hyperexcitability is a known driver of seizures[46], and our findings reveal hypoglycosylation of AT1B2 (Fig 3), a glycosylated component of the Na+/K+ pump, in ALG13-CDG. This hypoglycosylation may impair the pump’s ability to regulate membrane potential, increasing the likelihood of hyperexcitability. Additionally, our proteomics and scRNAseq analyses show upregulation of excitatory receptors such as *GRM1, GRIK2*, and *GRIA4*, further supporting a hyperexcitable neuronal environment.

#### Latent Hyperexcitability and Network Hypoactivity

Despite the molecular evidence of neuronal hyperexcitability, our electrophysiological recordings revealed network hypoactivity in ALG13-CDG hCOs, characterized by reduced firing rates, immature bursting patterns, and shorter neurite extensions. This paradox where hypoactive networks are present in a seizure-prone disease suggests the presence of latent hyperexcitability, a state in which neurons are biochemically primed to overactivate despite a subdued baseline functional state. Upregulation of excitatory receptors (e.g., GRM1, GRIA1) may represent an early response to suppressed activity (homeostatic plasticity), increasing the system’s vulnerability to overstimulation or destabilization.

Importantly, latent hyperexcitability has been reported in other neurodevelopmental and seizures disorders where stress can trigger a transition from hypoactivity to seizure[47–49]. This may explain why ALG13-CDG patients develop seizures despite early developmental suppression of neural activity. Furthermore, our pseudotime trajectory data revealed temporal mistiming in neuronal lineage development, with early inhibitory and delayed excitatory neuronal development. Such asynchronous development may impair proper excitatory/inhibitory (E/I) balance, creating an environment for seizures once excitatory circuits catch up in maturity.

Together, these findings highlight the importance of viewing ALG13-CDG and potentially other CDGs as disorders of network instability, where early hypoactivity drives downstream excitability changes that predispose to neurological pathology.

## Conclusion

While our study focuses on a cortical organoid model of ALG13-CDG, the molecular and functional mechanisms we identify—such as mistimed neuronal development, early network hypoactivity, and compensatory hyperexcitability—may reflect broader pathophysiological processes shared across CDGs with neurological involvement. Similar disruptions, including network hypoactivity and excitatory/inhibitory imbalance, have also been observed in 2D neuronal models of SLC35A2-CDG, highlighting common neurodevelopmental vulnerabilities. Together, our findings illustrate how multi-pathway dysregulation can converge to drive cortical dysfunction in CDG, providing a foundation for uncovering shared mechanisms and guiding the development of targeted therapeutic strategies.

## Acknowledgments

This work was supported by the 1U54NS115198-01 from the National Institute of Neurological Diseases and Stroke (NINDS), the National Center for Advancing Translational Sciences (NCATS), the National Institute of Child Health and Human Development (NICHD), and the Rare Disorders Consortium Disease Network (RDCRN). This work was also partially supported by a grant from DBT/Wellcome Trust India Alliance entitled “Center for Rare Disease Diagnosis, Research and Training” (IA/CRC/20/1/600002) to A.P. We thank Mayo Clinic DERIVE Office and Mayo Clinic Center for Biomedical Discovery for financial support and grants from NCI from (P30CA15083).

