## Supplementary Table, Method, and Figures for "ALG13 loss-of-function alters glycosylation, impairs neuronal maturation, and drives network hypoactivity in a cortical organoid model of CDG"

### Supplementary methods

#### Glycoproteomics LC-MS/MS

The peptides/glycopeptides were first trapped on a trap column (100 mm × 2 cm, Acclaim PepMap100 Nano-Trap, Thermo Fisher Scientific) at a flow rate of 20 μl/min before . LC separation was performed on an analytical column (EasySpray 75 μm x 50 cm, C18 2 μm, 100 Å) at a flow rate of 300 nl/min for 150 min using a linear gradient of 0.1% formic acid in water (solvent A) and 0.1% formic acid in acetonitrile (solvent B). All experiments were performed in a data-dependent acquisition (DDA) mode at an isolation window of 0.7 m/z. Data acquisition was performed with option of "lock mass" (m/z 441.12002) for all data.

#### QC, GEM formation, library preparation, and sequencing for scRNAseq

The cells were counted and measured for viability using the Vi-Cell XR Cell Viability Analyzer (Beckman-Coulter). All our samples passed this QC. At the same time, the GAC thawed barcoded Gel Beads from the -80 °C and prepared cDNA master mix according to the manufacture's instruction for Chromium Single Cell 3' Library and Gel Bead Kit (10x Genomics). Per sample the aim was to recover 10,000 cells and as such the cDNA master mix was added to the cells so that we have 500,000 cell per ml. Subsequently, these cells, Gel Beads, and partitioning oil were added to the Chromium Single Cell G chip. The filled chip was loaded into the Chromium X Controller, where each sample was processed and the individual cells within the sample were partitioned into uniquely labeled Gel Beads-In-Emulsion (GEMs). The GEMs were collected from the chip and taken to the bench for reverse transcription, GEM dissolution, and cDNA clean-up. The resulting cDNA contained a pool of uniquely barcoded molecules, and a portion of this cDNA was taken for library construction. To create gene expression libraries, standard Illumina sequencing primers and unique sample index (Dual Index Kit TT-Set A; 10x Genomics) were added to each cDNA pool. All cDNA pools and resulting libraries were quantified using Qubit High Sensitivity assays (Thermo Fisher Scientific) and Agilent Bioanalyzer High Sensitivity chips (Agilent). The libraries were subsequently sequenced at 50,000 fragment reads per cell by using the Illumina's standard protocol and the Illumina NovaSeq™ 6000 S4 flow cell, which reads 100 X 2 paired end reads using NovaSeq S4 sequencing kit and NovaSeq Control Software v1.8.0. Base-calling was performed using Illumina's RTA version 3.4.4.

***MALDI-QTOF Analysis of N-Glycans, processing and analysis***

Tissue samples were analyzed on a timsTOFfleX MALDI-QTOF mass spectrometer (Bruker) in positive mode using a 10 kHz SmartBeam 3D laser with a 20 µm spot size and raster, with 300 shots per pixel for high-resolution imaging of N-glycan spatial distribution (m/z 700-4000). MS data were processed in SCiLS Lab 2022b Pro, with N-glycan peaks manually selected based on theoretical mass values (±5 ppm) from an in-house database and prior MS/MS fragmentation data. Statistical significance was determined by Student t-test ($p < 0.05$). Metaboanalyst 5.0 generated PCA, heatmaps, and VIP score plots.

**X-inactivation**

Genomic DNA (gDNA) was isolated from cultured fibroblasts, induced pluripotent stem cells (iPSCs), and cultured cortical organoids using the QIAamp DNeasy Mini Kit (Qiagen). gDNA was digested with the DdeI restriction enzyme and the methylation-sensitive restriction enzyme HpaII (New England Biolabs) for 30 minutes at 37C, followed by 15 minutes heat inactivation at 80C. The c.56_287 portion of exon 1 of the androgen receptor (AR) gene on the X chromosome, which contains a polymorphic trinucleotide polyglutamine repeat region was amplified using the HotStarTaq DNA Polymerase kit (Qiagen). Amplicons were separated via gel electrophoresis and the relative abundance of the two polymorphic alleles were measured to assess X-inactivation skewing.

**scRNAseq integrated Seurat object generation**

The fastq files generated from sequencing were run through the 10X Genomics Cell Ranger pipeline to generate h5 files for each sample. Further analysis was conducted on these files using R studio and Seurat packages. The h5 files were merged and filtered to only keep cells that had RNA count greater than 800, feature RNA greater than 500, and mitochondrial gene percent less than 10%. After filtration, the data was normalized and integrated to eliminate batch effects and generate an integrated Seurat object via SelectIntegrationFeatures, FindIntegrationAnchors, and IntegrateData functions.

| Patient | CDG-0458 | CDG-1017 | CDG-11816 | CDG-11740 |
|---|---|---|---|---|
| Age at Time of Sample Collection | 2 Y | 9 Y | 3 Y | 6 Y |
| Genetic Variant | c.320A>G, N107S | c.320A>G, N107S | c.320A>G, N107S | c.320A>G, N107S |
| Developmental Delay | + | + | + | + |
| Seizures | + | + | + | + |
| Non-Verbal | + | + | + | + |
| Intellectual Disability | + | + | + | + |
| Central Hypotonia | - | + | + | - |
| Ophthalmological Abnormalities | - | - | + | + |
| Hearing Loss | - | - | - | + |
| ACTH therapy | + | + | + | + |
| Topamax trialed | +* | +* | - | + |
| Ketogenic diet trialed | + | + | + | + |

| | | | | |
|---|---|---|---|---|
| Ketogenic diet related seizures control | - | - | - | + |
| Seizure Control | - | + | - | + |

***Supplementary Table 1:*** *Clinical Data on the ALG13-CDG Patients in this Study*

Supplementary Figures

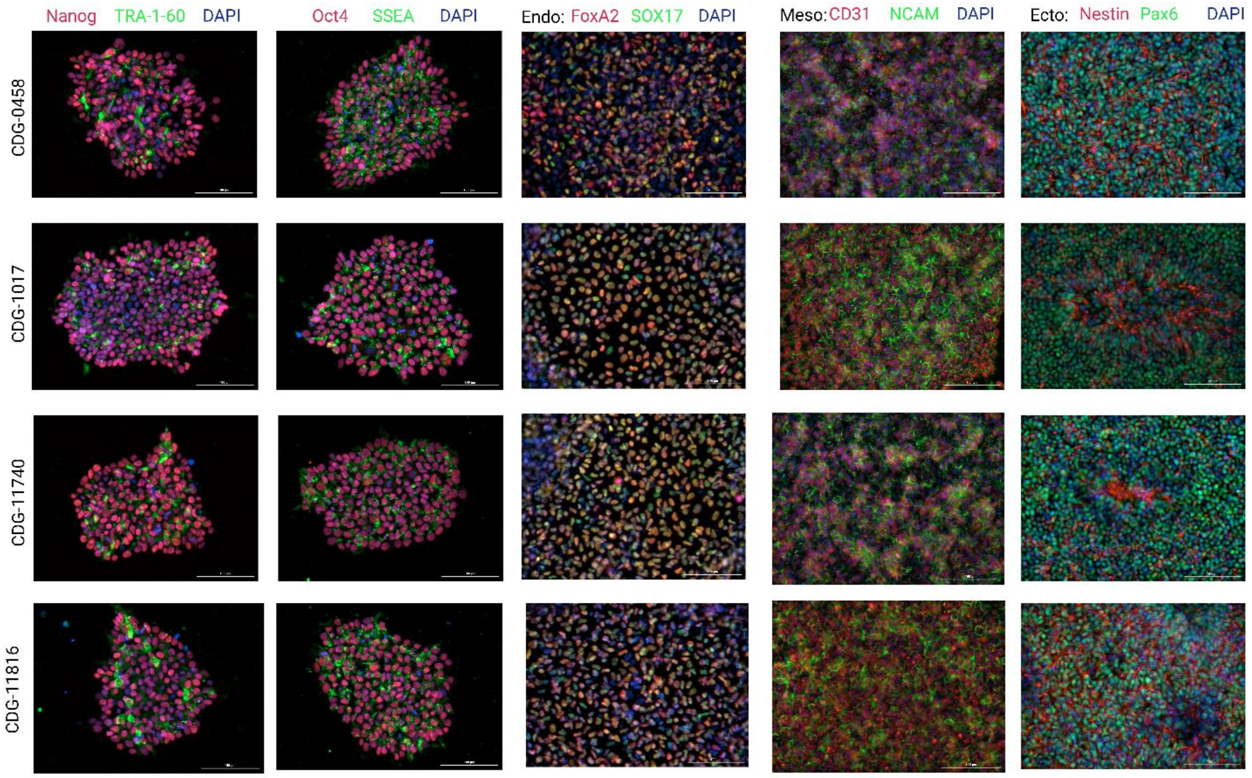


**Supplementary Figure 1: Pluripotency and Germ Layer Markers for ALG13-CDG iPSCs.** All images are at 20X resolution. Nanog, TRA-1-60, Oct4, & SSEA were used as pluripotency markers. SOX17 and FoxA2 staining was used as endoderm marker. CD31 and NCAM were used as mesoderm markers while Nestin and Pax6 were used as Ectoderm markers.

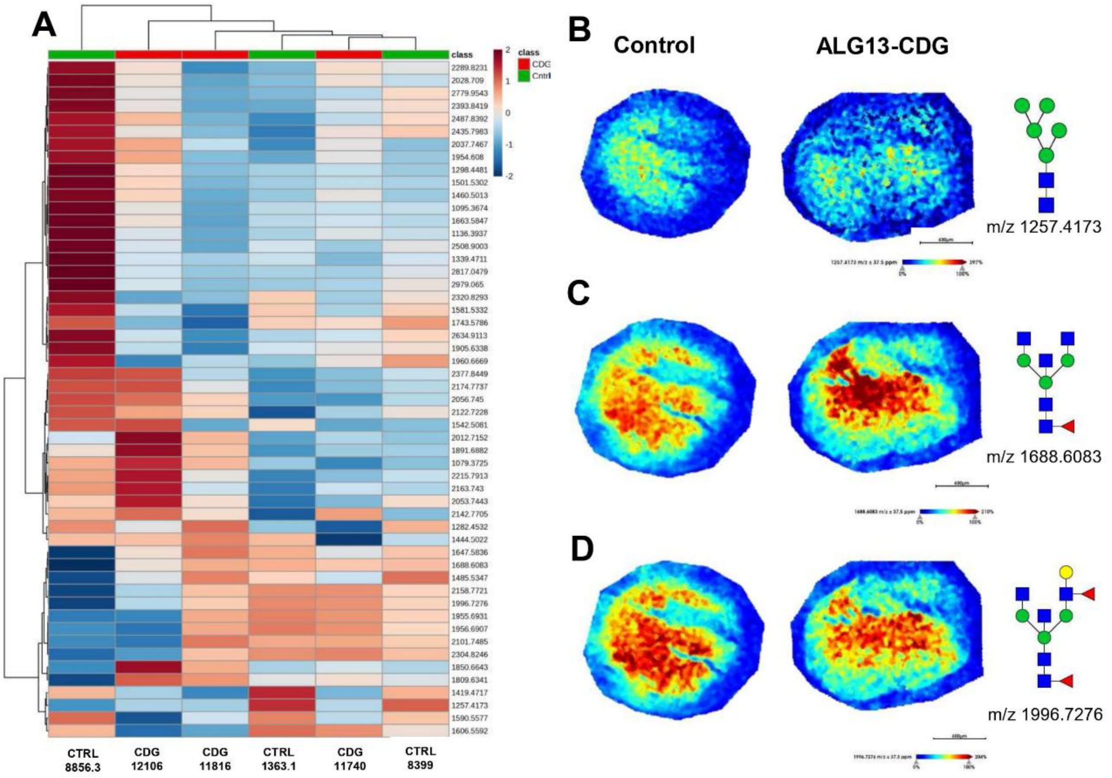


**Supplementary Figure 2 . N-glycomic comparison of ALG-13 deficient cortical organoidsto controls. A**) Unbiased clustering heatmap analysis of all 57 N-glycan features stratifying the groups. MALD-MSI representative images of m/z **B**)1257.4173 H5N2, **B**)1688.6083 H3N5F1, and **C**) 1996.7276 H4N5F2 N-glycans for control and ALG13 deficient cortical organoidswith corresponding relative abundance and N-glycan structure on the right. Intensity gradients from blue (least abundant) to red most abundant. Scale bars are below images.

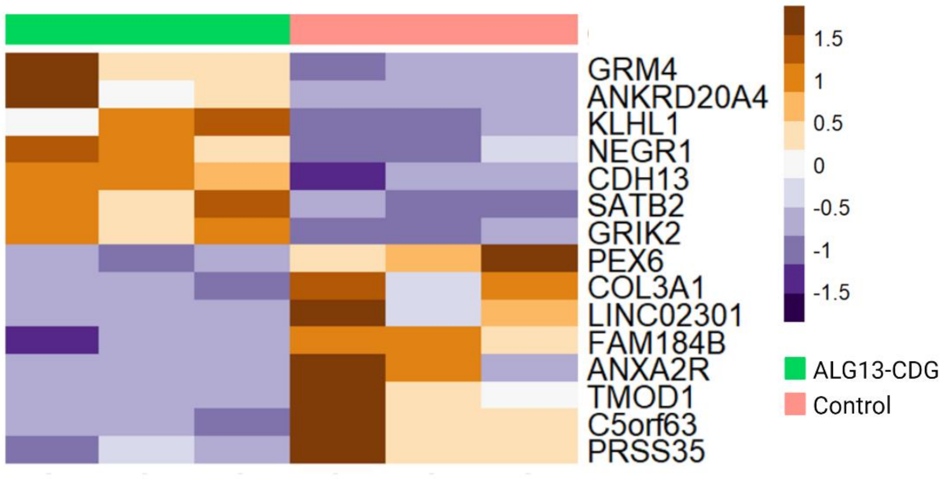


**Supplemntary Figure 3:Bulk differential gene expression analysis of scRNAseq data** reveals alteration in genes important for neruonal excitation, lipid metabolism, extracellular matrix, and lipid metabolism.

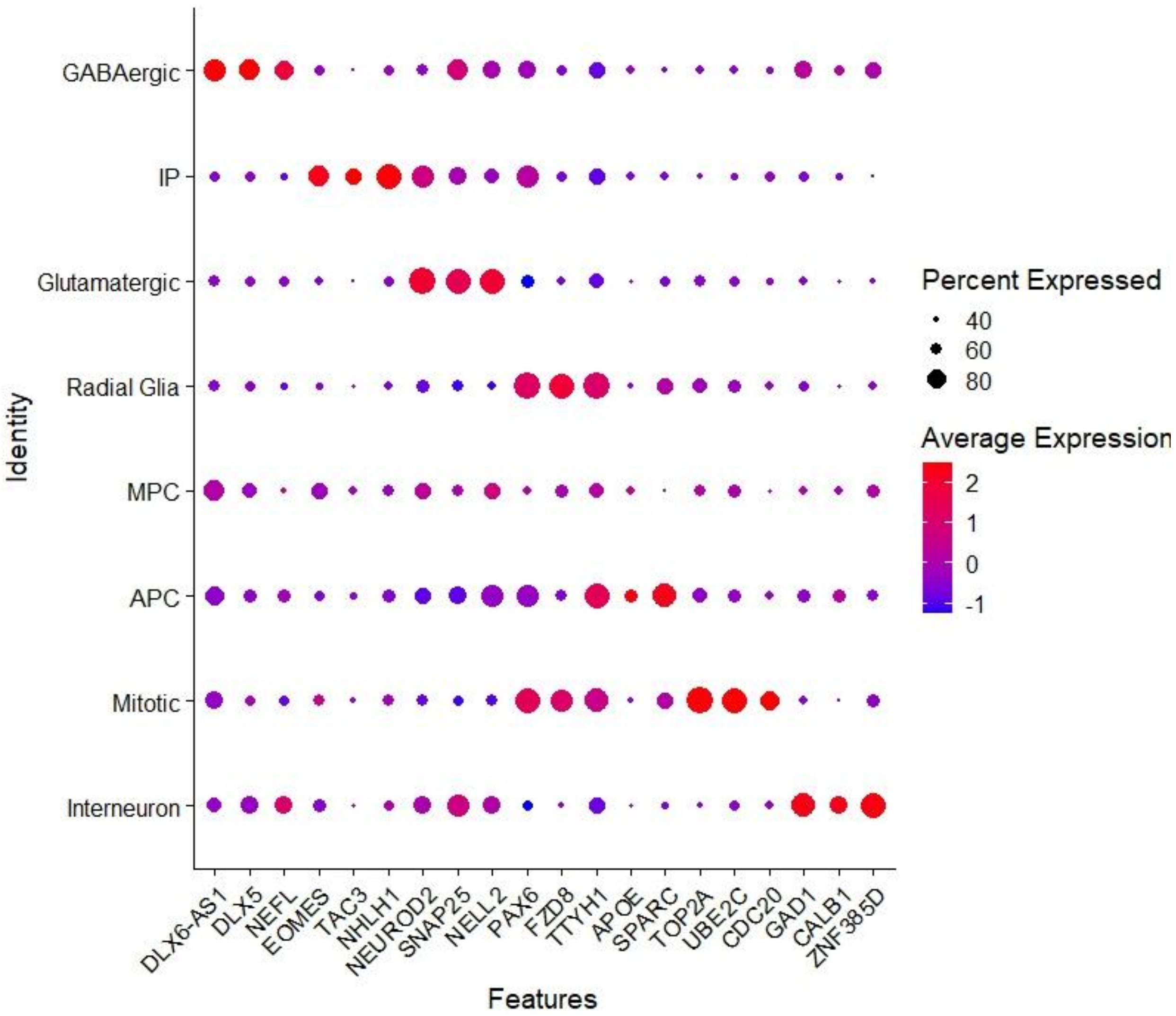


Supplementary Figure 4: Dotplot of transcripts associated with specific cell populations: Percent Expressed is represented by the size of the dot and expression level is represented on a scale from -1 (blue) to 2 (red).

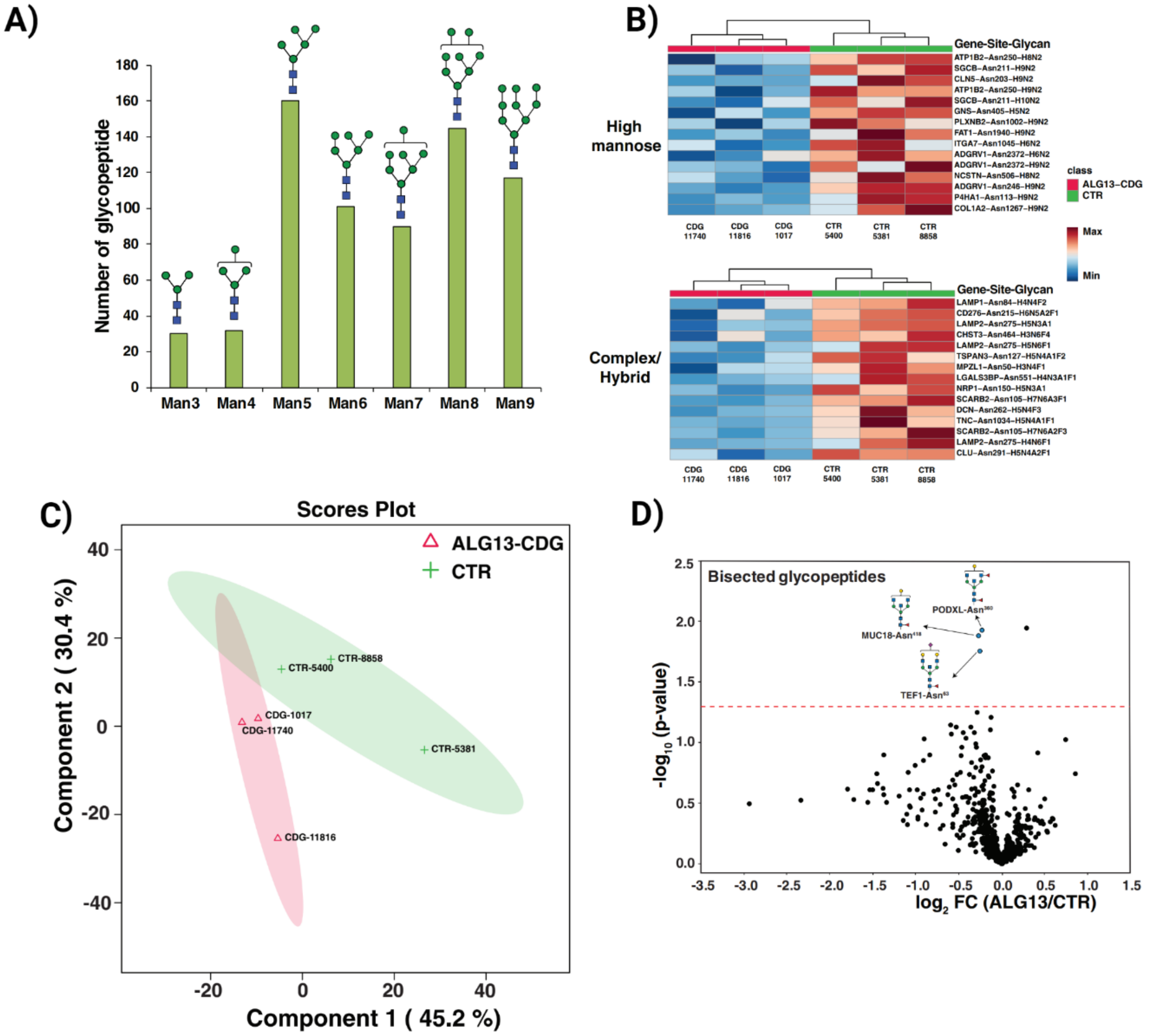


***Supplementary Figure 5. Site-specific glycosylation changes in patient derived ALG13-CDG hCOs.*** *(A) High mannose glycans containing glycopeptides are shown with numbers representing glycopeptides found with increasing hexose units. (B) Heatmap of significantly changing glycopeptides (p-value < 0.05) with different high-mannose and complex/ hybrid glycan moieties. The pattern is color coded. (C) Partial Least Squares Discriminant Analysis (PLS-DA) based on reporter ion intensities for all*

*identified glycopeptides of ALG13-CDG patients and controls. The percentage of total variance associated with each component is shown in brackets with the axis label. (D) Volcano plots depicting the downregulation of glycopeptides carrying the bisected glycan composition in ALG13-CDG. Putative structures are shown using Symbol Nomenclature for Glycans (SNFG).*

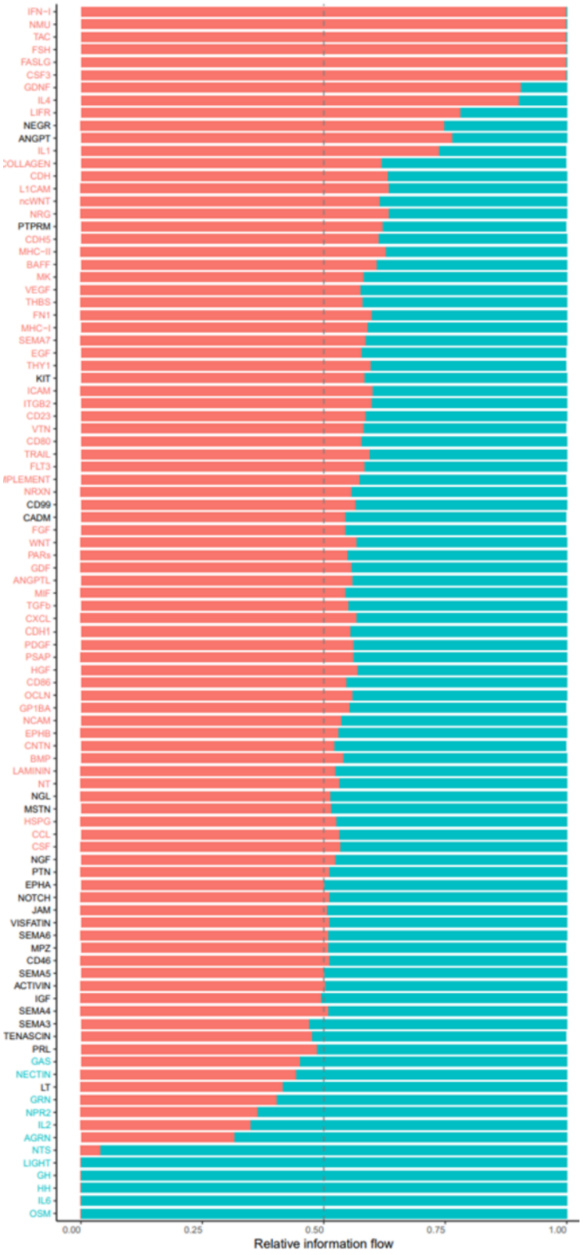


**Supplementary Figure 6**: Ranked flow chart showing the relative information flow from signaling pathways identified in hCOs. The cyan color indicates flow from controls and the punch pink color indicates flow from ALG13-CDG hCOs

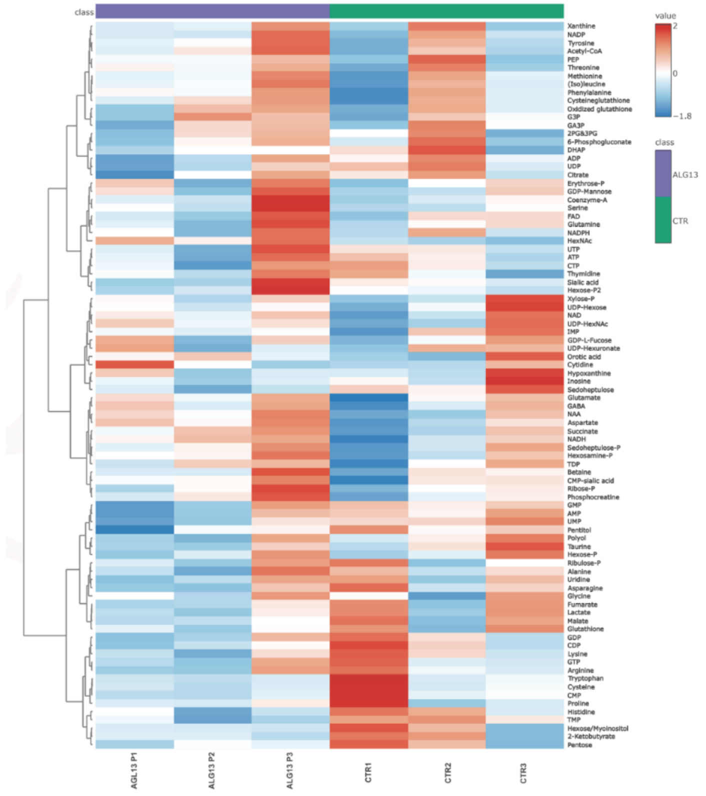


**Supplementary Figure 7**: *Heatmap of metabolites detected in ALG13 deficient and control hCOs. The pattern is color coded.*
